# Precision targeting of C3+ reactive astrocyte subpopulations with endogenous ADAR in an iPSC-derived model

**DOI:** 10.1101/2025.06.11.659213

**Authors:** Hyosung Kim, Andrew Kjar, Rebecca J. Embalabala, Jonathan M. Brunger, Ethan S. Lippmann

## Abstract

Astrocytes play pivotal roles in maintaining neural architecture and function. However, their pronounced heterogeneity, especially in reactive states where distinct subtypes can adopt potentially opposing functions (e.g., neuroprotective vs. neuroinflammatory), complicates our understanding of their net contributions to neurological disorders. A critical challenge arises because these functionally distinct subpopulations often coexist, and the lack of precise tools to separately monitor or manipulate them has significantly hindered efforts to dissect their specific roles in disease progression. Here, we address this gap by developing and optimizing fluorescent RNA sensors mediated by endogenous adenosine deaminase acting on RNA (ADAR) for application in induced pluripotent stem cell (iPSC)-derived astrocytes. We employed a streamlined screening methodology to enhance sensor specificity and functionality for complement component 3 (C3), a key marker predominantly associated with neuroinflammatory astrocytes, thus enabling subtype-specific tracking and providing a crucial tool for distinguishing these cells within heterogeneous populations. By integrating the biological complexity of astrocytes with the technological precision of ADAR-mediated sensing, this study establishes a robust framework for investigating astrocyte dynamics.

## Introduction

Astrocytes, once considered a homogeneous support cell population within the central nervous system (CNS), are now recognized for their substantial heterogeneity in both physiological and pathological contexts^1–5^. Under normal conditions, astrocytes exhibit region-specific molecular and functional characteristics. For example, cerebellar astrocytes are enriched in genes associated with synaptic transmission and plasticity, while striatal astrocytes prioritize the expression of metabolic genes^6^. Additionally, morphologically and molecularly distinct astrocytes are found across the layers of the cerebral cortex, further underscoring their specialized regional roles ^7^. The discovery of interlaminar astrocytes, a feature exclusive to primates, further emphasizes species-specific astrocytic diversity, suggesting that their specialization extends beyond regional to evolutionary dimensions^8–10^.

Pathologically, astrocyte heterogeneity is also pronounced, with distinct reactive phenotypes emerging in response to neurodegenerative diseases. For example, in Alzheimer’s disease (AD), astrocytes develop disease-associated phenotypes (DAAs) with unique transcriptional signatures, including upregulation of complement component 3 (C3), a key immune marker linked to neuroinflammation. C3 expression distinguishes DAAs from both homeostatic and other reactive astrocytes, suggesting a potential role in AD pathogenesis^11^. Similarly, in multiple sclerosis (MS), astrocytes exhibit heterogeneous responses: some adopt neuroinflammatory phenotypes, while others assume neuroprotective roles, potentially contributing to the variable disease outcomes observed in MS patients^11^. Such distinct and often opposing roles of astrocyte subtypes in pathological conditions highlight their critical importance for the development of therapeutic strategies. This underscores the need for advanced tools capable of precise identification and real-time characterization of astrocyte subtypes that will better enable cell type- specific, context-dependent monitoring and therapeutic approaches^12^.

A potential strategy for assessing astrocyte states in real-time is through the use of RNA sensors such as adenosine deaminases acting on RNA (ADAR), an enzyme that catalyzes the conversion of adenosine to inosine in double-stranded RNA. ADAR technology has shown particular promise in delineating cell- type-specific roles within complex tissues, such as the brain, where intricate interactions between diverse cell types influences both normal function and disease progression^13–17^. This capability is theoretically well-suited for targeting known astrocyte subtypes, as defined by transcriptional studies^18–23^ to enable direct investigation of their roles within neuroinflammatory processes. However, while ADAR-mediated sensors hold promise for this approach, their early-stage development and unresolved issues, including suboptimal efficiency and the absence of applications for iPSC-derived cells, highlight the need for further refinement. To address these limitations, we implemented systematic sequence optimization through screening-based approaches to enhance sensor specificity and functionality in human astrocytes derived from induced pluripotent stem cells (iPSCs), focusing solely on endogenous mechanisms without reliance on external elements beyond the sensor, thus improving monitoring accuracy and reliability. These improvements not only increase sensor accuracy but also broaden the applications of ADAR technology in both basic and translational research, offering a promising path for therapeutic strategies in neuroscience and cellular biology.

## Results

### ADAR-mediated RNA modification is effective in iPSCs

ADAR-based sensors represent a promising technology for detecting specific RNA molecules within cells. While their utility has been demonstrated effectively in various contexts and cell types, their application and systematic validation within human iPSCs—a critical model system for human development and disease—remain relatively unexplored. Therefore, to first establish feasibility, we sought to validate the functionality of a well-characterized ADAR sensor system in human iPSCs. To this end, we utilized a previously established sensor construct, expressing Blue Fluorescent Protein (BFP) constitutively and an enhanced green fluorescent protein (EGFP) reporter downstream of a sensor sequence containing a premature UAG stop codon (**Figure 1A**). Sensor activation, indicated by EGFP expression, requires ADAR-mediated editing of the UAG codon to UIG, which is dependent on the presence of a complementary trigger RNA. We co-transfected human iPSCs with the sensor plasmid and a separate plasmid expressing tdTomato as an exogenous trigger RNA. As a positive control, similar experiments were conducted in HEK293 cells, which have previously been utilized for this specific sensor^16^. In both cell types, EGFP expression was observed predominantly in cells co-expressing the tdTomato trigger, while cells lacking tdTomato showed negligible EGFP signal (**Figure 1B, D**). Quantitative analysis of fluorescence intensity in individual, doubly positive cells revealed a strong positive correlation between the tdTomato trigger signal and the EGFP reporter signal in both HEK293 cells (Pearson’s r = 0.86, **Figure 1C**) and iPSCs (Pearson’s r = 0.796, **Figure 1E**). These findings demonstrate that the ADAR-mediated RNA sensor system is functional in human iPSCs and that reporter activation correlates quantitatively with the levels of its target trigger tdTomato RNA.

**Figure 1.**
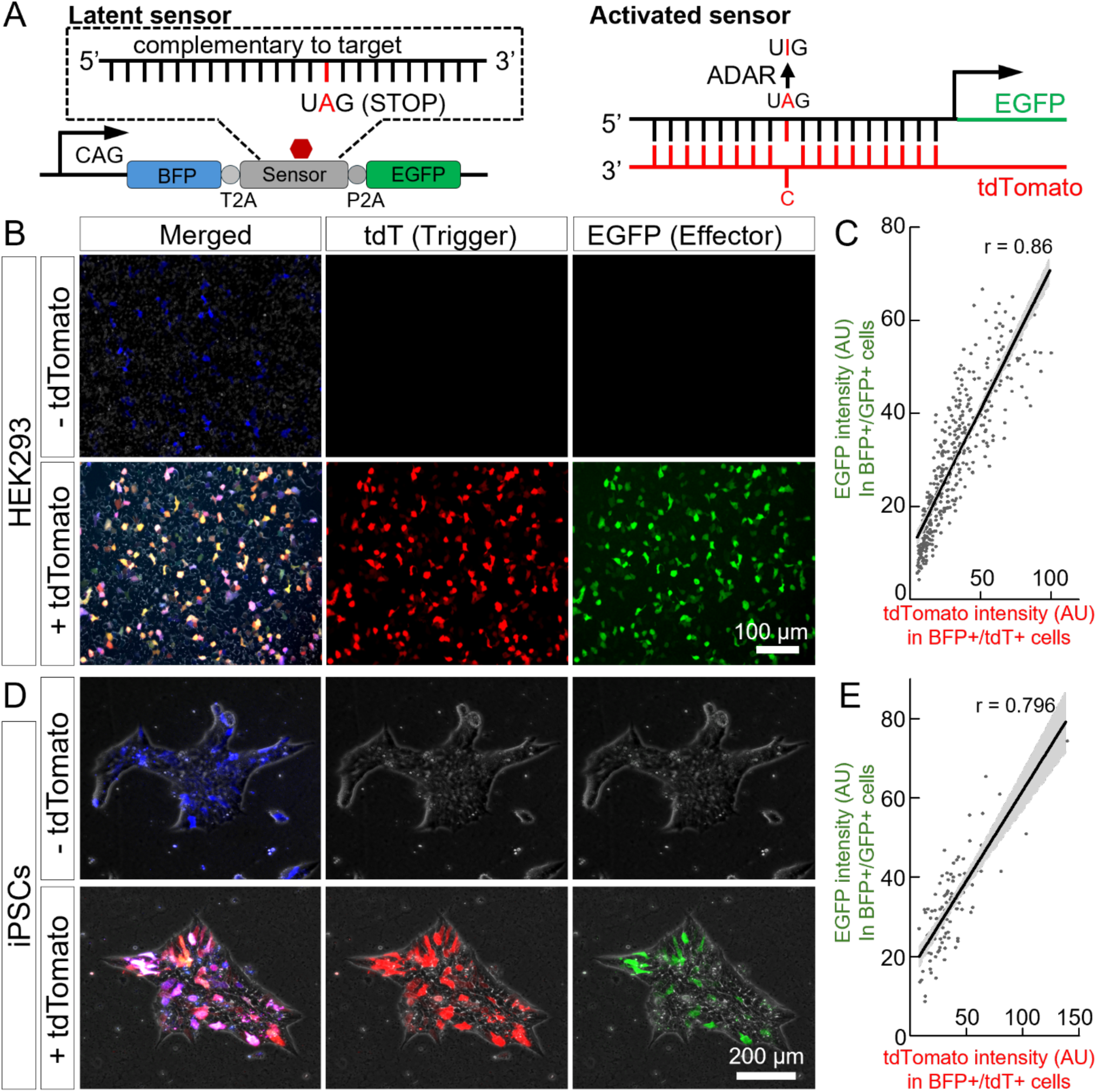
Validation of ADAR sensor system functionality in HEK293 cells and human iPSCs. **(A)** Schematic diagram illustrating the ADAR sensor mechanism. The sensor construct expresses BFP constitutively (driven by the CAG promoter). In the absence of the target RNA (tdTomato), a UAG stop codon within the sensor sequence prevents downstream EGFP translation (Latent sensor). Binding of the target RNA recruits endogenous ADAR enzymes, which edit the UAG stop codon to UIG, permitting EGFP translation (Activated sensor), between T2A and P2A allowing for multicistronic expression. **(B)** Representative fluorescence microscopy images of HEK293 cells transfected with the ADAR sensor construct (expressing BFP, blue) with (+) or without (-) co-transfection of the tdTomato trigger plasmid (tdT, red). Sensor activation is indicated by EGFP expression (green). **(C)** Representative scatter plot showing the correlation between tdTomato intensity (trigger level) and EGFP intensity (sensor activation) in individual BFP-positive HEK293 cells co-expressing tdTomato from one of three independent experiments. Each dot represents a single cell. Pearson’s correlation coefficient (r = 0.86) for the depicted experiment is indicated; similar positive correlation trends were confirmed across all three biological replicates. **(D)** Representative fluorescence microscopy images of human iPSCs transfected with the ADAR sensor construct (expressing BFP, blue/magenta) with (+) or without (-) co-transfection of the tdTomato trigger plasmid (tdT, red). Sensor activation is indicated by EGFP expression (green). **(E)** Representative scatter plot showing the correlation between tdTomato intensity (trigger level) and EGFP intensity (sensor activation) in individual BFP-positive iPSCs co-expressing tdTomato from one of three independent experiments. Each dot represents a single cell. Pearson’s correlation coefficient (r = 0.80) for the depicted experiment is indicated; similar positive correlation trends were confirmed across all three biological replicates. AU, arbitrary units.

### Human iPSC-derived astrocytes express ADAR and respond to ADAR-based RNA sensors

Given that the recruitment of endogenous ADAR is central to our strategy, we first examined existing single-cell RNA sequencing datasets for iPSC-derived neural cells^24^. This reanalysis revealed widespread ADAR transcript expression across the majority of neural cell types represented in the atlas, including various neuronal and glial lineages. Importantly, astrocytes displayed detectable ADAR expression at levels comparable to many other relevant cell types within this developmental context (**Figure 2A**). We next generated astrocytes from human iPSCs using an established multi-stage differentiation protocol carried out over 40 days (**Figure 2B**). The resulting cultures displayed characteristic astrocyte morphology and expressed the canonical astrocyte markers GFAP and CD44, confirming successful differentiation (**Figure 2C**). To determine if the ADAR sensor system validated in undifferentiated iPSCs (**Figure 1**) was also functional in this differentiated cell type, we co-transfected the iPSC-derived astrocytes with the BFP- sensor plasmid and the tdTomato trigger plasmid. Consistent with previous observations, EGFP reporter expression, indicating sensor activation, was induced specifically in astrocytes co-expressing the tdTomato trigger RNA (**Figure 2D**). Furthermore, quantitative analysis of fluorescence intensities in individual cells revealed a significant positive correlation between the tdTomato trigger levels and EGFP reporter activation (Pearson’s r = 0.612, **Figure 2E**). These results confirm that iPSC-derived astrocytes express sufficient endogenous ADAR and possess the necessary cellular machinery to support the function of the ADAR-based RNA sensor system, with reporter output correlating with target RNA abundance.

**Figure 2.**
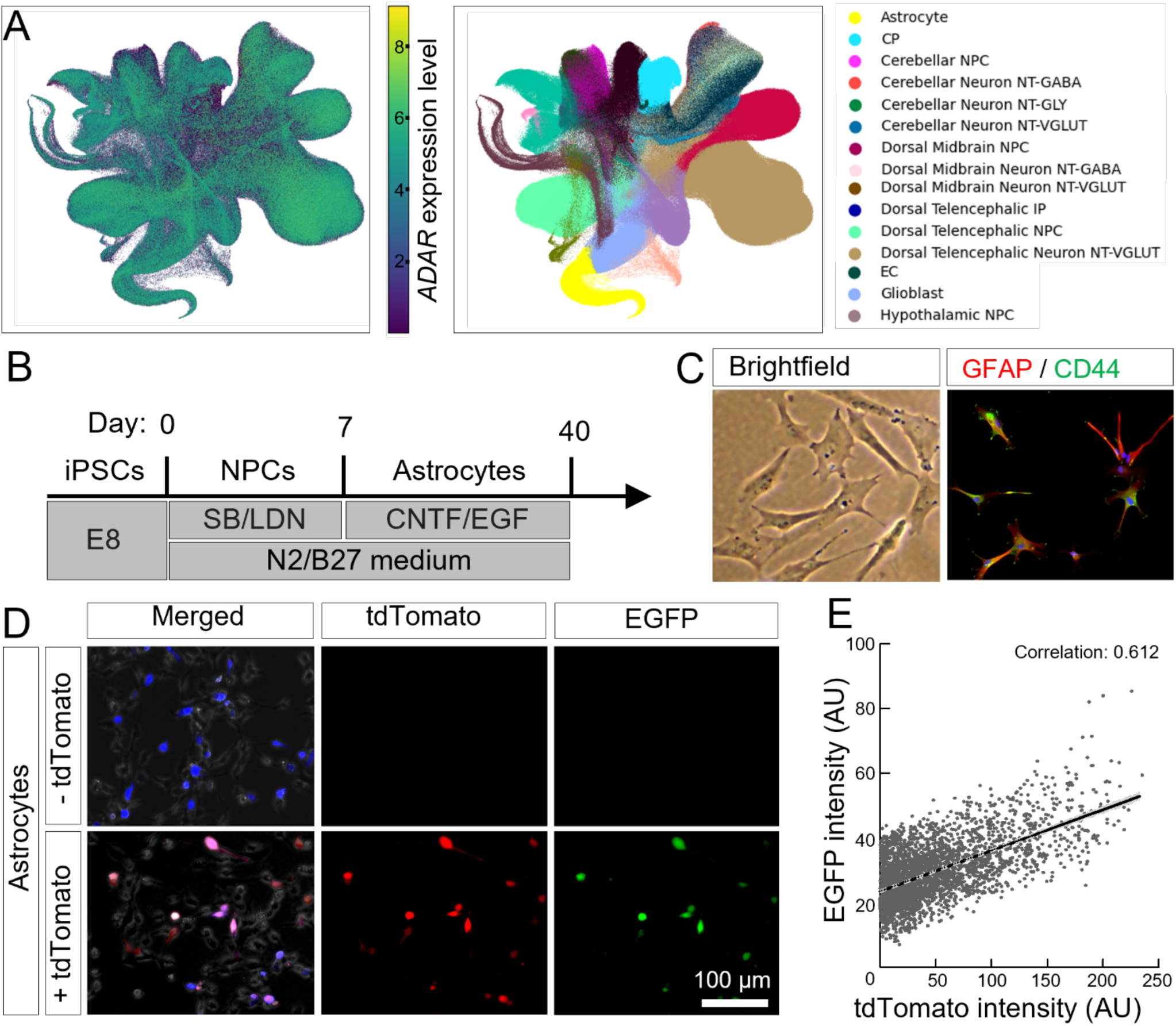
ADAR sensor system functions in iPSC-derived astrocytes. **(A)** Re-analysis of single-cell RNA sequencing data from a reference human neural organoid atlas^24^. UMAP plots show ADAR expression levels (left panel; color scale indicates expression level from low (blue) to high (yellow)) and clustering of annotated cell types (right panel; key provided), including various neuronal and glial populations. Astrocytes (yellow) display notable ADAR expression relative to several other cell types shown. **(B)** Schematic overview of the differentiation protocol used to generate astrocytes from human iPSCs over 40 days. Key stages (iPSCs, NPCs, Astrocytes), media components (E8, SB/LDN, CNTF/EGF, N2/B27), and time points are indicated. **(C)** Representative brightfield image (left) and immunofluorescence staining (right) for astrocyte markers GFAP (red) and CD44 (green) after day 40 of differentiation. **(D)** Representative fluorescence microscopy images of astrocytes transfected with the sensor construct expressing the constitutive marker BFP, with (+) or without (-) co-transfection of the tdTomato trigger plasmid. Images show tdTomato (red) and EGFP (green, indicating sensor activation). **(E)** Representative scatter plot showing the correlation between tdTomato intensity (trigger level) and EGFP intensity (sensor activation) in individual marker-positive astrocytes co-expressing tdTomato from one of three independent experiments. Each dot represents a single cell. Pearson’s correlation coefficient (r = 0.612) for the depicted experiment is indicated; similar positive correlation trends were confirmed across all three biological replicates. AU, arbitrary units.

### Screening identifies effective ADAR sensors for endogenous C3 mRNA in iPSC-derived astrocytes

Having established that the ADAR sensor system functions in iPSCs and iPSC-derived astrocytes with an exogenous trigger (**Figure 1**, **Figure 2**), we next sought to develop sensors capable of detecting an endogenous mRNA target relevant to astrocyte reactivity. We focused on C3, as our previous research indicated that C3 expression is negligible in quiescent astrocytes but is highly upregulated by inflammatory cytokines^25,26^, making it an ideal target for sensing specific reactive states. To identify optimal sensor sequences, we synthesized 20 distinct sensor candidates targeting various regions within the C3 mRNA, including sequences with deliberate single or double mismatches intended to potentially modulate editing efficiency or specificity. Each sensor candidate, along with an mCherry marker, was cloned into a PiggyBac transposon vector for stable genomic integration (**Figure 3A**). Pools of iPSCs stably expressing each sensor construct were generated and subsequently differentiated into astrocytes (**Figure 3B**). To induce endogenous C3 expression, these astrocyte cultures were treated with a combination of TNFα, IL-1α, and C1q (T.I.C.), which are cytokines known to promote an inflammatory reactive astrocyte phenotype associated with C3 upregulation. Bulk RNA-sequencing analysis confirmed a marked increase in C3 transcript levels in astrocytes following T.I.C. treatment compared to non-treated controls (**Figure 3C**). We then evaluated the performance of the 20 sensor candidates by quantifying the ADAR-mediated UAG-to-UGG editing efficiency at the target site within the sensor transcript via RNA sequencing in both non-treated and T.I.C.-treated conditions (**Figure 3D**). The screen revealed a range of activities among the candidates; several showed minimal or no editing, while others exhibited varying degrees of basal editing in the non-treated state. Notably, a subset of sensors, including candidates #2, #14, and #15, demonstrated a substantial increase in editing efficiency specifically upon T.I.C. treatment, correlating with the induced C3 expression (**Figure 3D**). This screening process successfully identified promising guide sequences for the robust ADAR-mediated sensing of endogenous C3 mRNA in iPSC-derived astrocytes.

**Figure 3.**
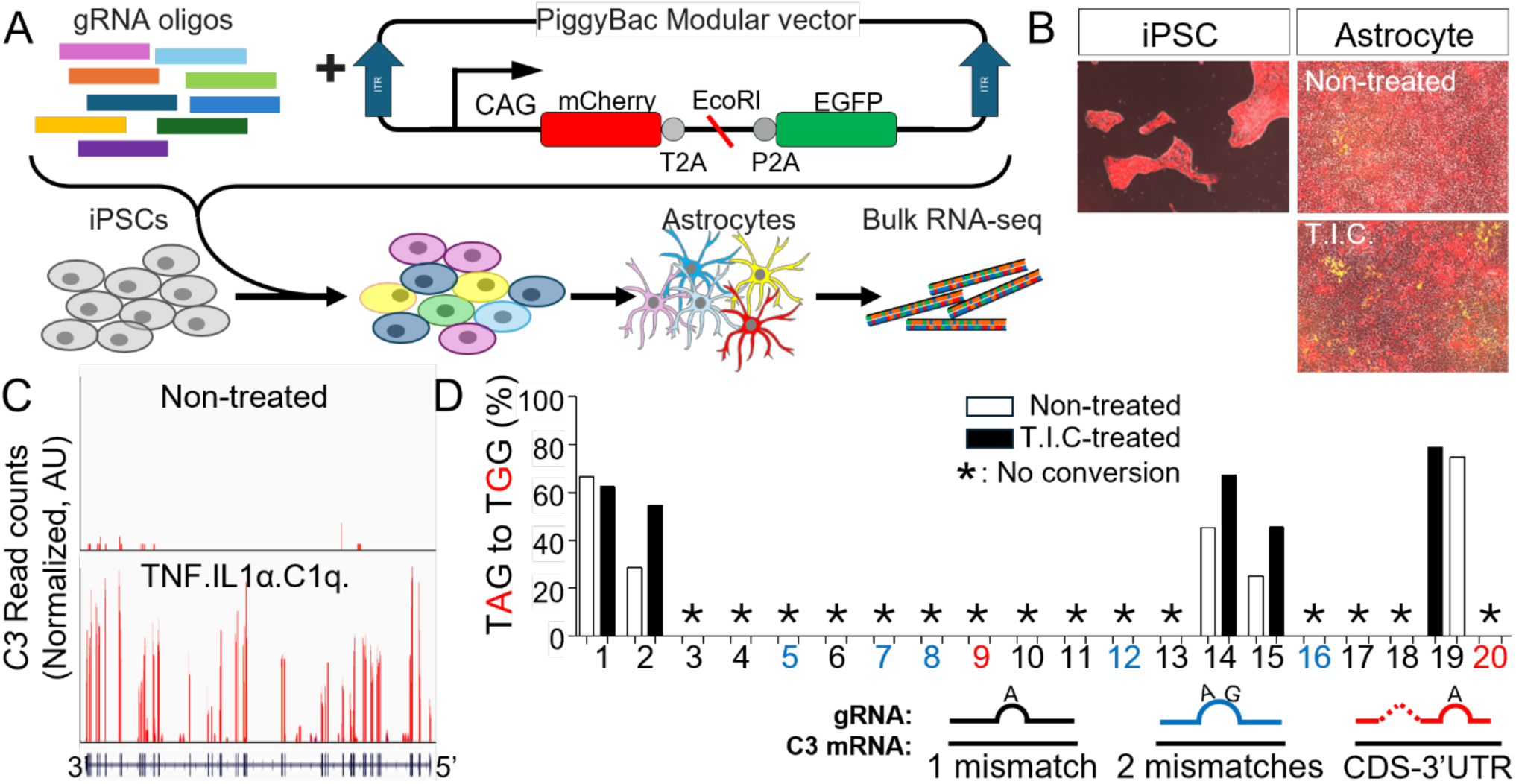
Screening of ADAR sensors targeting endogenous C3 mRNA. **(A)** Schematic illustrating the sensor screening strategy. Twenty distinct gRNA sensor sequences targeting C3 were cloned into a PiggyBac modular vector expressing mCherry and EGFP under a CAG promoter, separated by T2A and P2A sequences. iPSCs were transfected with the pooled sensor library, stably integrated using PiggyBac transposase, and differentiated into astrocytes. Bulk RNA-sequencing was then performed on astrocyte populations. **(B)** Representative images showing mCherry expression overlaid on phase contrast images in the stable iPSC line (left panel) and derived astrocytes under non-treated (top right panel) or T.I.C.-treated (bottom right panel) conditions. **(C)** Representative RNA-sequencing read coverage plots across the C3 gene locus in non-treated (top) and T.I.C.-treated (TNFα, IL-1α, C1q; bottom) astrocytes, demonstrating induction of C3 expression upon T.I.C. treatment. Normalized read counts (AU., arbitrary units) are shown on the y-axis, and the C3 transcript structure (exons as boxes, introns as lines) is shown below. **(D)** Quantification of UAG-to-UGG editing efficiency (%) for each of the 20 C3 sensor candidates via bulk RNA-sequencing in non-treated (white bars) and T.I.C.-treated (black bars) astrocytes. Sensor candidates targeted the C3 3’UTR and incorporated 1 or 2 mismatches as indicated by the key below the graph. Asterisks (*) denote candidates with no detectable editing conversion. Data are derived from one non-treated biological replicate and two T.I.C.-treated biological replicates.

### Lentiviral delivery of a selected C3 sensor enables fluorescence-based detection of endogenous C3 induction

Following the initial screen that identified promising C3 sensor candidates based on RNA editing efficiency (**Figure 3**), we selected a lead candidate sensor (#2) for further validation using a different delivery method and readout. Lentiviral transduction was chosen to achieve stable sensor expression and overcome the typically low efficiency observed with non-viral transfection methods (e.g., lipofection) in differentiated astrocytes. To assess sensor performance at the single-cell level via fluorescence microscopy using this stable expression approach, we packaged the selected C3 sensor sequence into a lentiviral vector co-expressing mCherry as a transduction marker (**Figure 4A**). Human iPSC-derived astrocytes were transduced with this lentivirus, with mCherry expression confirming successful transduction (**Figure 4A**, micrograph). To induce endogenous C3 expression, sensor-equipped astrocyte cultures were treated with the pro-inflammatory cytokines TNF. qRT-PCR analysis confirmed that TNF treatment significantly upregulated C3 mRNA expression relative to GAPDH in these astrocytes (**Figure 4C**, left panel; P=0.0003). Sensor activation was subsequently evaluated by measuring EGFP reporter fluorescence intensity in individual mCherry-positive cells (**Figure 4B**). Compared to non-treated control cells which displayed minimal EGFP signal, astrocytes treated with TNF exhibited a significant increase in EGFP fluorescence intensity (**Figure 4C**, right panel; P<0.001). These results validate that the selected C3 sensor, delivered via lentivirus, can effectively and sensitively report the upregulation of its endogenous target mRNA, C3, in response to an inflammatory stimulus using a fluorescent output detectable at the single-cell level.

**Figure 4.**
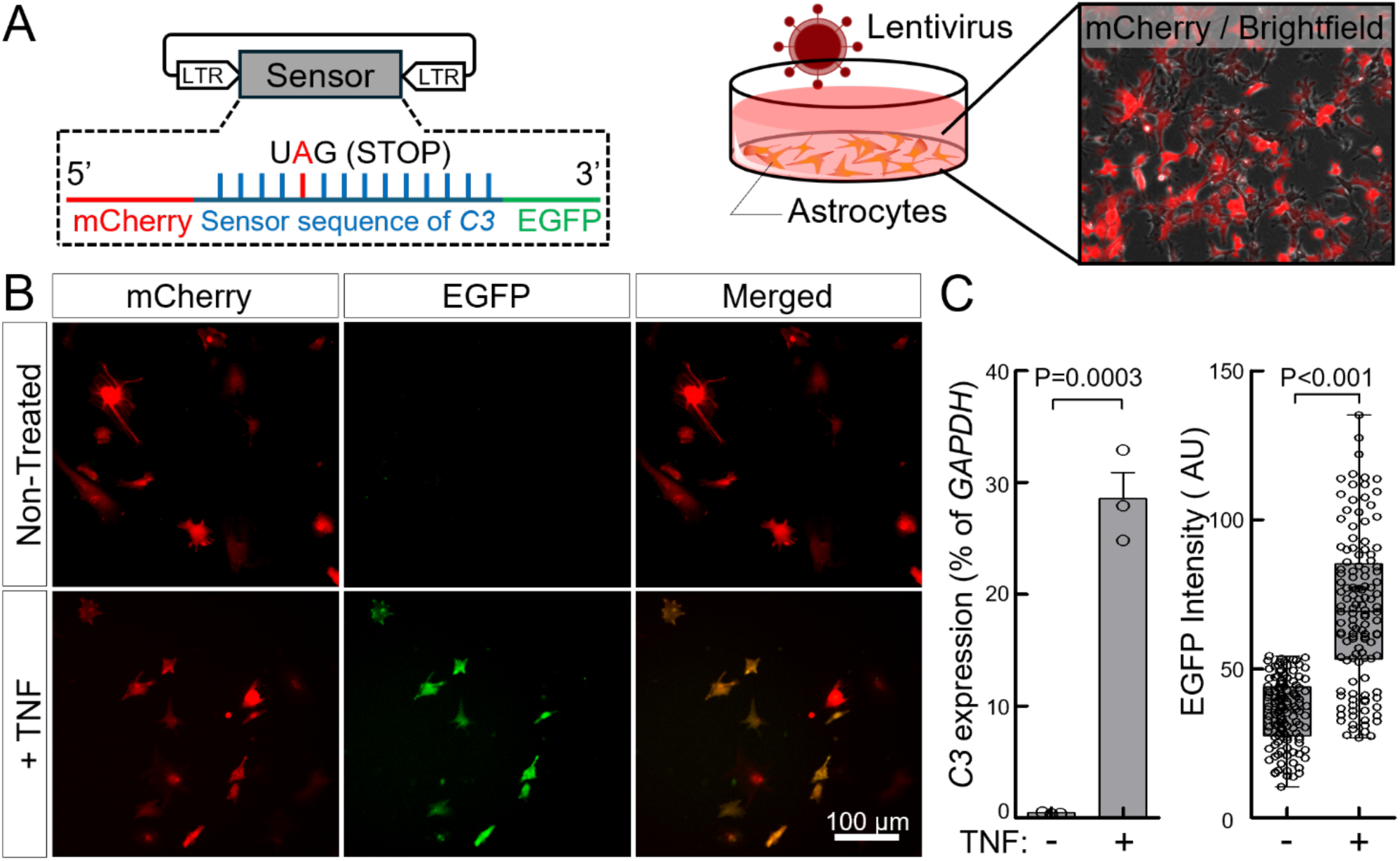
Lentiviral C3 sensor reports endogenous C3 induction in iPSC-derived astrocytes. **(A)** Schematic of the experimental approach for validating the selected C3 sensor (#2) using lentiviral delivery. Astrocytes were transduced with a lentivirus encoding the sensor construct (right panel shows construct details: LTRs, CMV-intron promoter driving mCherry, P2A, C3 (#2) sensor sequence, T2A, EGFP with UAG stop codon). Representative overlay of mCherry fluorescence and brightfield image shows transduced astrocytes (middle panel). **(B)** Representative fluorescence microscopy images of lentivirus-transduced astrocytes treated without (Non-Treated) or with TNF (+TNF) for 48 hours. Images show mCherry (indicating transduced cells) and EGFP (indicating sensor activation). **(C)** Quantitative PCR analysis of *C3* mRNA expression (left panel) in iPSC-derived astrocytes that are untreated (-) or TNF-treated (+) for 48 hours. *C3* mRNA expression is shown as a percentage relative to *GAPDH* housekeeping gene expression. Data are presented as mean ± SEM from three independent experiments. Statistical significance (P=0.0003) was determined using an unpaired two-tailed Student’s t-test. Quantification of EGFP fluorescence intensity (right panel) in individual mCherry-positive cells under non-treated (-) and TNF-treated (+) conditions. Data are presented as box plots showing median, quartiles, and whiskers, with individual cell measurements overlaid. Statistical significance was determined using an unpaired two-tailed Student’s t-test (GraphPad Prism) (P<0.001). Data are representative of three independent experiments, which showed similar trends. AU, arbitrary units.

## Discussion

In this study, we developed and validated an RNA sensor framework leveraging endogenous ADAR activity to identify and monitor C3-expressing astrocytes, a subpopulation associated with neuroinflammatory conditions, within human iPSC-derived models. This approach circumvents the need for exogenous enzyme overexpression, potentially minimizing global off-target editing and enhancing biological relevance, particularly within the complex context of iPSC-derived neural cultures. Crucially, by directly sensing the presence of the target C3 mRNA, this system offers distinct advantages over traditional promoter-based reporter constructs. These systems primarily reflect the activity of a specific gene’s regulatory elements and transcriptional initiation, which may not always directly correlate with the abundance of the mature, functional transcript due to post-transcriptional regulation or variations in promoter activity across different cellular states or differentiation stages. In contrast, our ADAR-based sensor directly interrogates the presence of the target RNA molecule itself, potentially offering a more immediate readout of transcript availability. Our findings demonstrate that this endogenous ADAR-mediated system can effectively report on target RNA presence, responding dynamically to neuroinflammatory stimuli in a subgroup of reactive astrocyte. This work provides a refined methodology for dissecting astrocyte heterogeneity with high precision, offering valuable tools for investigating specific cellular subpopulations in models of neurological disease.

The successful application of this endogenous ADAR strategy underscores its potential for distinguishing subpopulation-specific markers within complex neural cultures. We initially confirmed the system’s basic functionality and quantitative response using an exogenous tdTomato trigger, observing strong correlations between trigger levels and sensor output in both undifferentiated iPSCs and iPSC-derived astrocytes. Crucially, the system proved effective for detecting an endogenous target; following T.I.C.- induced C3 upregulation, a screen of 20 sensor candidates identified several sequences exhibiting robust UAG-to-UGG editing increases. Further validation using lentiviral delivery of a lead sensor confirmed a significant increase in fluorescence upon TNF treatment, demonstrating reliable reporting at the single-cell level. These results support the hypothesis that endogenous ADAR activity, when coupled with optimized sensor sequences, provides a sensitive means to track dynamic changes in astrocyte state, such as the induction of C3 in response to inflammatory cues.

The implications of these findings are substantial for studying astrocyte biology, particularly the challenge of dissecting the roles of heterogeneous subpopulations with potentially opposing functions in neurological disorders. By enabling the specific tracking of C3-positive astrocytes, this tool provides a means to isolate and study this specific subpopulation, contributing to a clearer understanding of its contribution to neuroinflammation distinct from other astrocyte subtypes (e.g., potentially neuroprotective populations). Practically, this approach offers a robust method for monitoring astrocyte reactivity in real- time within human cell models. This RNA-editing framework could be valuable in drug discovery, facilitating screens for compounds that selectively modulate C3+ astrocyte populations, and holds promise for translational studies investigating astrocyte subtype roles in neurodegenerative conditions.

A key advantage of the ADAR sensor platform described herein lies in its intrinsic adaptability to diverse biological contexts, an attribute enabled by our integrated high-throughput screening pipeline. Translation of this technology to novel systems—such as primary astrocyte cultures, in vivo models, or alternative mRNA targets—requires strategies capable of addressing the inherent biological variability that typifies complex tissues. Variables such as signaling network heterogeneity, differentiation state, local RNA secondary structure, and cell type-specific expression patterns of ADAR isoforms or RNA-binding proteins can markedly influence sensor efficacy. To overcome this context-dependency, we developed a robust and scalable empirical screening framework that obviates the need for predictive modeling. This platform enables rapid identification of optimally functional sensor sequences within the specific cellular environment of interest. By directly assaying performance in situ, this approach ensures compatibility across distinct signaling milieus, cell types, and potentially even across species. Thus, while the use of endogenous ADAR enzymes circumvents issues related to ectopic enzyme activity and the design of guide sequences confers target specificity, it is the integration of our context-sensitive screening strategy that endows the system with its versatility. This modular, adaptable framework positions the ADAR sensor platform as a broadly applicable tool for customized RNA-based applications across diverse biological systems.

Future studies should focus on validating this ADAR sensor system in primary human astrocytes and potentially in vivo models to confirm its utility beyond iPSC systems. Expanding the investigation to include other inflammatory stimuli and astrocyte subtype markers would broaden the applicability of this approach. Exploring the integration of this sensor technology with complementary methods, such as CRISPR-based gene activation or inhibition targeting specific astrocyte subtypes, could provide powerful synergistic tools for functional studies. Moreover, systematically investigating the factors contributing to cell-line variability in sensor performance will be important for standardizing this technology for wider use in diverse experimental contexts and disease models.

## Methods

### iPSC maintenance

The CC3 iPSC line was used for all experiments^27^. Undifferentiated iPSCs were maintained in E8 medium on 6-well plates coated with growth factor reduced Matrigel (Corning). iPSCs were passaged with Versene (Thermo Fisher Scientific) upon reaching 60-80% confluency.

### iPSC differentiation to astrocytes

iPSCs were dissociated using Accutase and plated onto Matrigel-coated plates at a density of 4x10^5^ cells/well in E8 medium supplemented with 10 µM Y-27632 (ROCK inhibitor, Tocris). On Day 0, the medium was switched to initiate neural induction using DMEM/F12 supplemented with N2 and B27 supplement (Gibco) and dual SMAD inhibitors (10 µM SB431542, Tocris; 100 nM LDN193189, Tocris). This neural induction medium (N2/B27 + SB/LDN) was replaced daily. On Day 7, the medium was changed to astrocyte differentiation medium consisting of DMEM/F12 supplemented with N2, B27, 10 ng/mL CNTF (Peprotech), and 10 ng/mL EGF (Peprotech). This medium was refreshed every 48 hours, and cells were passaged upon reaching approximately 80% confluency, until Day 40.

### Cytokine treatments on astrocytes

For inflammatory astrocytic activation, astrocyte cultures at 80% confluence in 6-well plates (in E6 medium containing CNTF and EGF) were treated with vehicle control or a cytokine cocktail (T.I.C.) consisting of 20 ng/mL TNF (R&D Systems), 10 ng/mL IL-1α (Thermo Fisher), and 500 ng/mL C1q (MyBioSource) for 48 hours. In specific cases, astrocytes were treated solely with TNF.

### C3 sensor candidates and PiggyBac modular vector construction

The PiggyBac transposon modular vector was designed using Benchling software and synthesized by VectorBuilder Inc. The vector contained the following key elements in order: an EF1A promoter driving mCherry (constitutive marker), a P2A sequence, an EcoRI cloning site for sensor insertion, a T2A sequence, and EGFP (sensor effector module). Additionally, the vector included a Hygromycin resistance gene driven by a CMV promoter for selection purposes. Twenty distinct sensor candidate sequences (72 or 90 bp) targeting the human C3 3’UTR were designed based on information from https://rnasensing.bio/^15^, incorporating either 1 or 2 mismatches. These sequences were synthesized as gBlocks (Integrated DNA Technologies, IDT) and their identity is listed in Supplemental Table 1. For cloning into the EcoRI-linearized PiggyBac vector using NEBuilder HiFi DNA Assembly (New England Biolabs), 15 bp flanking sequences homologous to the vector ends were added to each gBlock oligo during synthesis. Successful insertion of sensor sequences was initially verified by PCR amplification across the cloning site followed by agarose gel electrophoresis to confirm expected band size. The presence and sequence of the sensor transcripts were further confirmed during the analysis of bulk RNA-sequencing data (see Bioinformatic Analysis section).

### Lentiviral C3 sensor vector construction

To construct the lentiviral C3 sensor vector, the sensor cassette containing P2A, the selected C3 sensor sequence C3-2, T2A, and EGFP was excised from the PiggyBac C3 sensor vector using BbvCI and BtsI restriction enzymes. This fragment was cloned into a lentiviral backbone vector (synthesized by VectorBuilder Inc.) linearized with EcoRI and MscI, which contained an mCherry marker driven by a CMV-intron promoter. Assembly was performed using NEBuilder HiFi DNA Assembly (New England Biolabs), utilizing 15 bp overlaps homologous to the linearized vector ends and the excised sensor cassette fragment, following the manufacturer’s protocol. The final construct links the CMV intron-driven mCherry to the sensor-EGFP cassette via the P2A sequence.

### Stable iPSC line generation by PiggyBac transposon vector

For generating stable iPSC lines expressing C3 sensors, iPSCs were co-transfected with a pooled library containing all 20 C3 sensor PiggyBac plasmids and the pEF-1α-HA-m7pB transposase expression vector (a gift from Dr. Lauren Woodard, Vanderbilt University Medical Center)^28^ using Lipofectamine 3000 Reagent (Invitrogen) according to the manufacturer’s recommended protocol and plasmid ratio. One day post-transfection, selection was initiated by adding Hygromycin at a standard concentration of 100 µg/mL to the culture medium. Selection was maintained for 7 days. Pools of stably transfected, hygromycin-resistant cells were subsequently expanded for differentiation experiments.

### Lentivirus production and transduction

Lentivirus was produced by co-transfecting HEK293T cells with the lentiviral C3 sensor transfer vector, a packaging plasmid (psPAX2, Addgene #12260), and an envelope plasmid (pMD2.G, Addgene #12259) using Lipofectamine 3000. Viral supernatant was collected at 48 and 72 hours post-transfection, filtered through a 0.22 µm filter, and concentrated using Lenti-X Concentrator (Takara Bio). For transduction, concentrated lentiviral particles produced from one 35 mm dish of HEK293T cells were resuspended in astrocyte medium and added to iPSC-derived astrocytes plated at a density of 100,000 cells/cm². Cells were incubated with the virus for 48 hours without polybrene, after which the medium was replaced with fresh astrocyte medium.

### qPCR analysis

Total RNA was extracted with TRIzol and converted to cDNA using a Superscript III First-Strand kit (Invitrogen). qPCR was performed on a BioRad CFX96 Thermocycler with TaqMan primers against *C3* and *GAPDH* (Thermo Fisher Scientific). Samples were analyzed by normalizing expression levels to *GAPDH* and relative quantification was performed using the standard 2-ΔΔC_t_ method.

### Immunostaining

Cells were fixed with 4% paraformaldehyde (PFA) in PBS, permeabilized with 0.3% Triton X-100 in PBS, and blocked with 10% goat serum in PBS. Primary antibodies were diluted in blocking solution and incubated overnight at 4°C. The following primary antibodies were used: chicken anti-GFAP (Aves, SKU: GFAP, 1:300) and mouse anti-CD44 (Abcam, AB6124, 1:1000 dilution). After washing with PBS, cells were incubated with corresponding Alexa Fluor-conjugated secondary antibodies (donkey anti- chicken/anti-mouse, Alexa Fluor 488, 647, Thermo Fisher Scientific, 1:1000 dilution) for 1 hour at room temperature. Nuclei were counterstained with DAPI for 10 min. Coverslips were mounted using Fluoromount-G (Southern Biotechnology).

### Microscopy and image analysis

Fluorescence images were acquired using a Leica DMi8 epifluorescence microscope equipped with 10x, 20x, 40x, and 60x objectives and appropriate filter sets. Images for quantification were primarily acquired using the 10x and 20x objectives. Image acquisition parameters were kept consistent across conditions for comparison. For quantitative analysis of fluorescence intensity, images were analyzed using ImageJ software (v1.53c). In select cases, customized scripts utilizing ImageJ macro functions were employed for analysis (code located in Supplemental Information). Individual cells were identified and segmented based on BFP and mCherry fluorescence. The mean fluorescence intensity of EGFP or tdTomato within each segmented cell was measured after background subtraction.

### RNA extraction and sequencing

Total RNA was extracted from astrocyte cultures (non-treated and T.I.C.-treated) using TRI Reagent (Fisher Scientific) according to the manufacturer’s instructions. Isolated RNA was further purified using an RNeasy Plus Mini Kit (Qiagen). RNA concentration was assessed using a NanoDrop spectrophotometer. Only samples with an RNA Integrity Number (RIN) value above 9.0 proceeded to library preparation. Purified RNA samples were submitted to the Vanderbilt Technologies for Advanced Genomics (VANTAGE) core facility, where RNA quality was further validated prior to library preparation and sequencing. Following ribosome depletion, cDNA libraries were prepared using the TruSeq RNA Sample Prep Kit (Illumina). Sequencing was performed on an Illumina NovaSeq 6000 platform, generating 150 bp paired-end reads. Raw sequencing data were received as paired-end, unmerged FASTQ files.

### Bioinformatic analysis

Reads were aligned to the human reference genome (hg19, GRCh37) using HISAT2 (v2.2.1), generating SAM alignment files. These SAM files were converted to BAM format, sorted, and indexed using Samtools (v1.21). Alignment quality and coverage, particularly over the C3 locus, were optionally visualized using the Integrative Genomics Viewer (IGV, v.2.19.3). Gene expression counts, particularly for C3 exons and introns, were quantified from the sorted BAM files using featureCounts (Subread package v2.1.0). Raw FASTQ files were used directly for editing efficiency analysis. Reads containing the sensor sequence were identified and extracted using custom Python scripts designed to recognize the flanking T2A and P2A sequences. Within these filtered reads, A-to-I (UAG-to-UGG or A-to-G at the target adenosine) editing efficiency was further quantified using custom Python code, available at https://github.com/LippmannLab/RNA-seq/blob/main/RNA%20sensor_Kim%20et.%20al. Editing percentage was calculated for each sensor in both non-treated and T.I.C.-treated conditions.

## Data availability

The raw RNA sequencing data have been submitted to ArrayExpress (E-MTAB-15239).

## Acknowledgments

This work was supported by NIH grant RF1 129735 to E.S.L. Pilot funding to H.K. was provided by the Vanderbilt Institute for Clinical and Translational Research (VICTR), which was supported by NIH CTSA award UL1 TR002243. A.K. was supported by the National Science Foundation Graduate Research Fellowship Program. RNA sequencing was performed by the Vanderbilt Technologies for Advanced Genomics (VANTAGE) core facility, which is supported in part by NIH grants 5UL1 RR024975, P30 CA068485, P30 EY008126, UL1 TR002243, and G20 RR030956. We are grateful to Dr. Lauren Woodard (Vanderbilt University Medical Center) for providing the pEF-1α-HA-m7pB transposase expression vector. We also thank members of the Lippmann laboratory for insightful discussions and critical feedback on the manuscript.

## Author Contributions Statement

H.K. conceived the study, designed the methodology, performed the experiments and formal analysis, created the visualizations, and wrote the original draft of the manuscript. A.K. contributed to the investigation and validation, including the reanalysis of transcriptomic data from iPSC-derived neural cells. R.E. conducted investigations and provided key resources, including the iPSC-derived astrocytes used in the study. J.M.B. assisted with methodology and investigation by providing resources and expertise for cloning and participated in scientific discussions. E.S.L. co-conceived the study, supervised the project, acquired funding, and reviewed and edited the manuscript. All authors have read and approved the final manuscript.

