## Supplemental information for "Precision targeting of C3+ reactive astrocyte subpopulations with endogenous ADAR in an iPSC-derived model"

### Supplemental Table 1. Sensor sequences screened in this study.

| | |
|---|---|
| C3-1 | GGCGTGGTGTGGGCGTGGCGCAGGGGCGTGACAATGGTGTGGGCGTAGCATGGGCGGGGCAGTCGGGCGGTCGCGCGCACGCGCAGGGAA |
| C3-2 | GGTGTGGGCGTGGCGCAGGGGCGTGACAATGGTGTGGGCGTGGCATAGGCGGGGCAGTCGGGCGGTCGCGCGCACGCGCAGGGAAAGCAC |
| C3-3 | GTGAAGGCGCCGAGGTCCTGGCATTGTTTCTGGTTCTCTTCGTCTTAGCATTCGTCCTCCTCGGGCCAGTGCTCCACCCAAGTGTCCTTC |
| C3-4 | ACAACCATGCTCTCGGTGAAGGCGCCGAGGTCCTGGCATTGTTTCTAGTTCTCTTCGTCTTGGCATTCGTCCTCCTCGGGCCAGTGCTCC |
| C3-5 | GGGGGTGTGGTCAGTTGGGGCACCCAAAGACAACCATGCTCTCGGTAGAGGCGCCGAGGTCCTGGCATTGTTTCTGGTTCTCTTCGTCTT |
| C3-6 | GGGCACCCAAAGACAACCATGCTCTCGGTGAAGGCGCCGAGGTCCTAGCATTGTTTCTGGTTCTCTTCGTCTTGGCATTCGTCCTCCTCG |
| C3-7 | GTGGGAACAGGTGAGGTTTCAAGTAGGATGGAGCTGAGCTGCAGGTAGGGCGGCTGGGGATTTCAGCCTCTCCCTCTTGGCAAAGAACTC |
| C3-8 | GAGAAAATGCGGTGGGAACAGGTGAGGTTTCAAGTAGGATGGAGCTAGGCTGCAGGTGAGGCGGCTGGGGATTTCAGCCTCTCCCTCTTG |
| C3-9 | CATGCTCTCGGTGAAGGCGCCGAGGTCCTGGCATTGTTTCTGGTTCTCTTCGTCGGAAGGGGCGTGGTGTAGGCGTGGCGCAGGGGCGTG |
| C3-10 | GGTGAAGGCGCCGAGGTCCTGGCATTGTTTCTGGTTCTCTTCGTCTAGGCATTCGTCCTCCTCGGGCCAGTGCTCCACCCAAGTGTCCTT |
| C3-11 | CCCAAAGACAACCATGCTCTCGGTGAAGGCGCCGAGGTCCTGGCATAGTTTCTGGTTCTCTTCGTCTTGGCATTCGTCCTCCTCGGGCCA |
| C3-12 | GCTGCAGGTGAGGCGGCTGGGGATTTCAGCCTCTCCCTCTTGGCAAAGAACTCCCACGTGAGATATAACTAGAGCTTTATCTGGAGTGGG |
| C3-13 | GTGGCGCAGGGGCGTGACAATGGTGTGGGCGTGGCATAGGCGGGGCAGTCGGGCGGTCGCGCGCACGCGCAG |
| C3-14 | TGGGCGTGGCGCAGGGGCGTGACAATGGTGTGGGCGTAGCATGGGCGGGGCAGTCGGGCGGTCGCGCGCACG |
| C3-15 | AAGAATGGCAGGTCAGGAAGGGGCGTGGTGTGGGCGTAGCGCAGGGGCGTGACAATGGTGTGGGCGTGGCAT |
| C3-16 | GGTGAGGTTTCAAGTAGGATGGAGCTGAGCTGCAGGTAGGGCGGCTGGGGATTTCAGCCTCTCCCTCTTGGC |
| C3-17 | CTCTCGGTGAAGGCGCCGAGGTCCTGGCATTGTTTCTAGTTCTCTTCGTCTTGGCATTCGTCCTCCTCGGGC |
| C3-18 | CCGAGGTCCTGGCATTGTTTCTGGTTCTCTTCGTCTTAGCATTCGTCCTCCTCGGGCCAGTGCTCCACCCAA |
| C3-19 | AAGACAACCATGCTCTCGGTGAAGGCGCCGAGGTCCTAGCATTGTTTCTGGTTCTCTTCGTCTTGGCATTCG |
| C3-20 | ACACTAGCAGGCGAACGCCAGGAGAAAATGCGGTGGGAACAGGTGAGGTTTCAATTCAGCCTCTCCCTCTAGGCAAAGAACTCCAGACAC |

**Supplementary Code**

```
// ImageJ Macro for Batch Channel Quantification
// This macro is a general-purpose example for batch processing multi-channel images.
// It first asks the user to specify which channel to quantify. Then, for each image,
// it isolates that channel, applies an automatic threshold, and measures fluorescence intensity.
// Users should adjust thresholding methods based on their specific imaging data.

macro "Generic Batch Channel Quantification" {

    // --- Create a dialog box for user input ---
    Dialog.create("Quantification Settings");
    Dialog.addNumber("Enter channel to quantify (e.g., 1-4):", 3); // Set a default value, e.g., 3
    Dialog.show();
    channelToQuantify = Dialog.getNumber();

    // 1. Setup for batch processing_Select a directory
    dir = getDirectory("Choose a Source Directory");
    list = getFileList(dir);
    setBatchMode(true);
    run("Clear Results");

    // 2. Loop through all files in the directory
    for (i=0; i<list.length; i++) {
        path = dir+list[i];
        open(path);
        imgName = getTitle();

        // 3. Isolate the user-defined channel of interest
        run("Split Channels");
```

```
        targetChannelName = "C" + channelToQuantify + "-" + imgName;

        // Close all other channels automatically
        for (c=1; c<=4; c++) {
            if (c != channelToQuantify) {
                selectWindow("C" + c + "-" + imgName);
                close();
            }
        }

        // 4. Select the target channel and proceed with measurement
        selectWindow(targetChannelName);

        // Define which parameters to measure
        run("Set Measurements...", "area mean standard redirect=None decimal=3");

        // Apply a threshold to identify objects (cells)
        // NOTE: This method should be adjusted by the end-user for optimal results.
        setAutoThreshold("Otsu dark");

        // Perform the measurement
        run("Measure");
        close();
    }

    // 5. Display the final results table
    setBatchMode(false);
    selectWindow("Results");
}
```
